## Supplementary Figures and Tables for "Impact of Rap-Phr system abundance on adaptation of *Bacillus subtilis*"

* Shared first authors

[#] corresponding author

**Present address:**

[6] Department of Biology, Memorial University of Newfoundland, St. John’s, NL, Canada

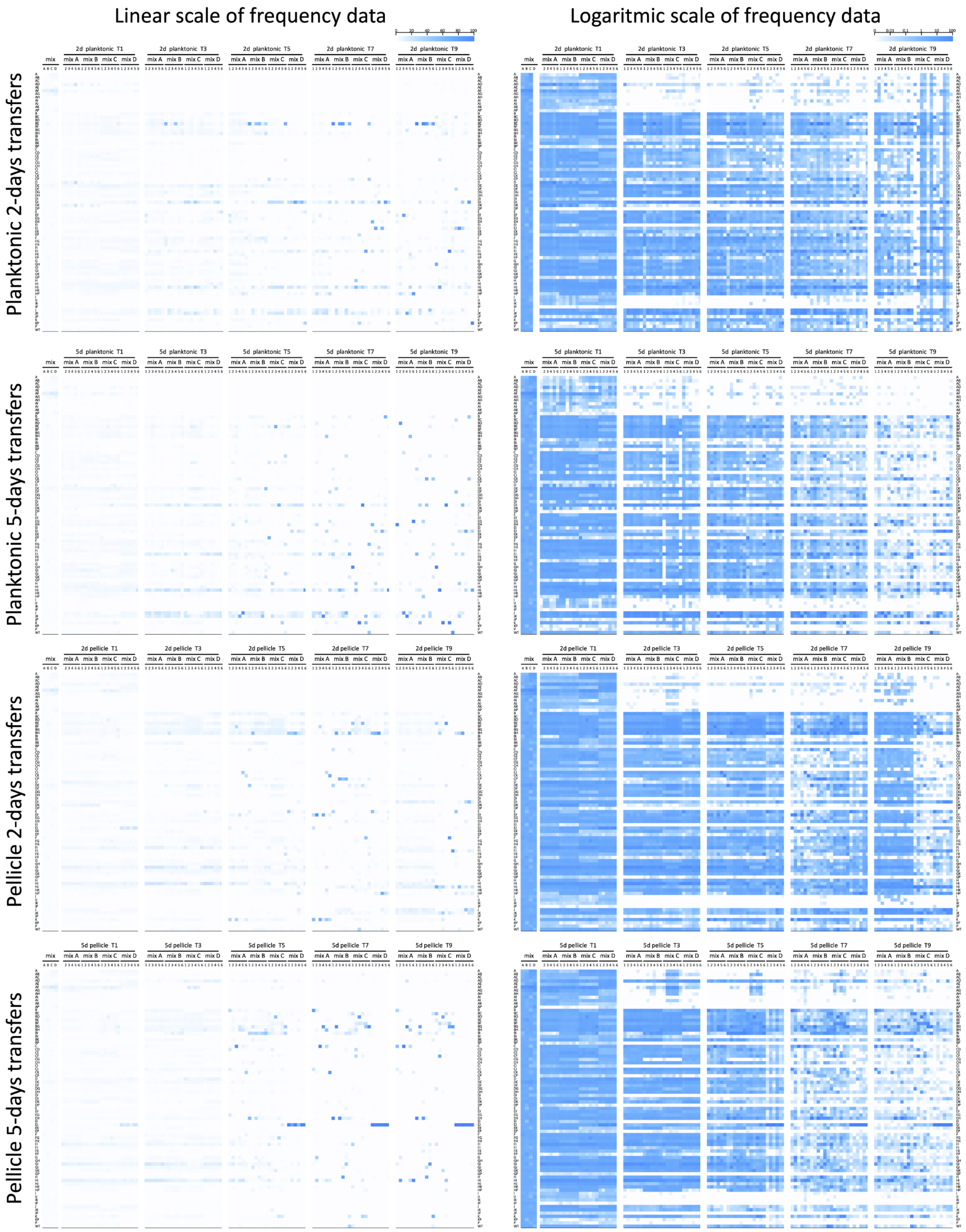


**Supplementary Fig. 1 Heat map representation of the population dynamics of *B. subtilis rap-phr* mutants in competition.** Boxes represent the population percentage of strains. Text columns at far-left and far-right indicate which *rap-phr* genes have been deleted (A indicates a Δ*rapA* mutant, AB indicates a Δ*rapA*Δ*rapB* mutant, and so on), WT indicates *B. subtilis* DK1042. Text rows on top indicate type of culture (planktonic or pellicle), incubation period (2d= 2 days, 5d= 5days), transfer number of represented population (t1, t3, t5, t7, and t9), and mix and replicate number. Competition populations were started from 4 population mixes (A to D), with 6 replicates per mix. The first two box columns indicate the population representation of tested strains in the competition starter mixes. Left and right panels show the same data, although using different scales (shown in the top-right corner of each panel): linear increment of percentage (left) and logarithmic increment of percentage (right).

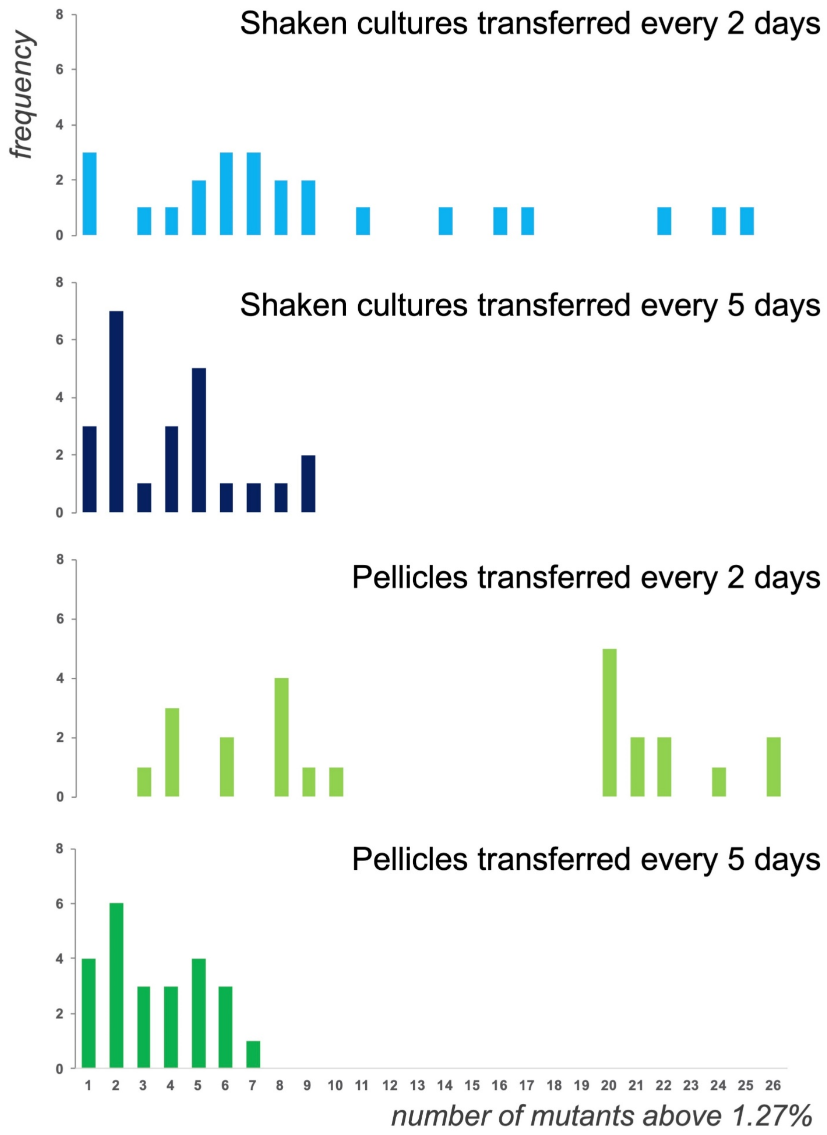


**Supplementary Fig. 2 Distribution within the 24 replicates of the number of mutants showing higher relative abundance than the initial 1.27% input, after transfer 9.** The X axis of the bar charts represents the number of mutants above 1.27% relative abundance at the end of the competition experiments of the 4 cultivation conditions, while y axis depicts their frequencies within the 24 replicates.

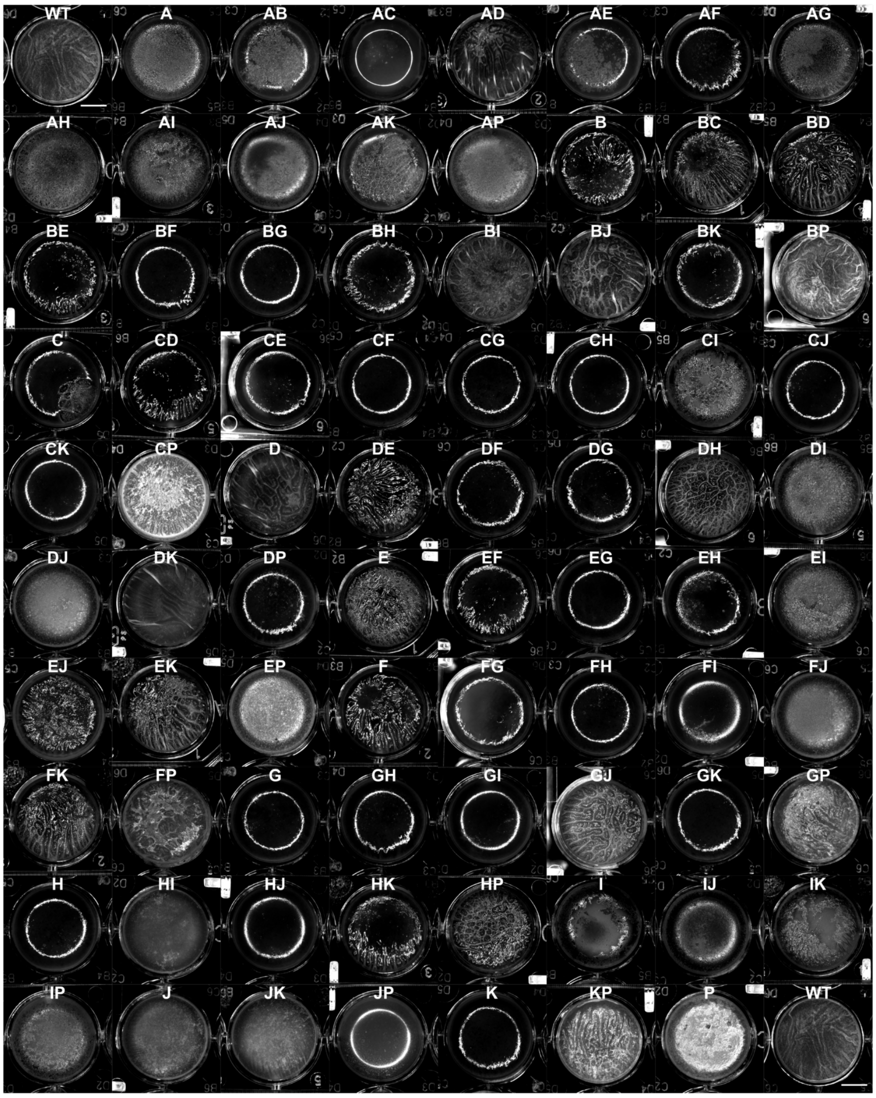


**Supplementary Fig. 3. 2 days pellicles of *B. subtilis* DK1042 and *rap-phr* mutants.** Bright-field images are shown after 2 days of incubation on MSgg medium at 30°C. WT indicates *B. subtilis* DK1042, A indicates a Δ*rapA* mutant, AB indicates a Δ*rapA*Δ*rapB* mutant, and so on. The images shown here have an adjusted contrast, so that the pellicles can be easily appreciated. The same image of *B. subtilis* DK1042 is presented twice (top-left and bottom-right) to facilitate pellicle comparison. The scale bars represent 5 mm.

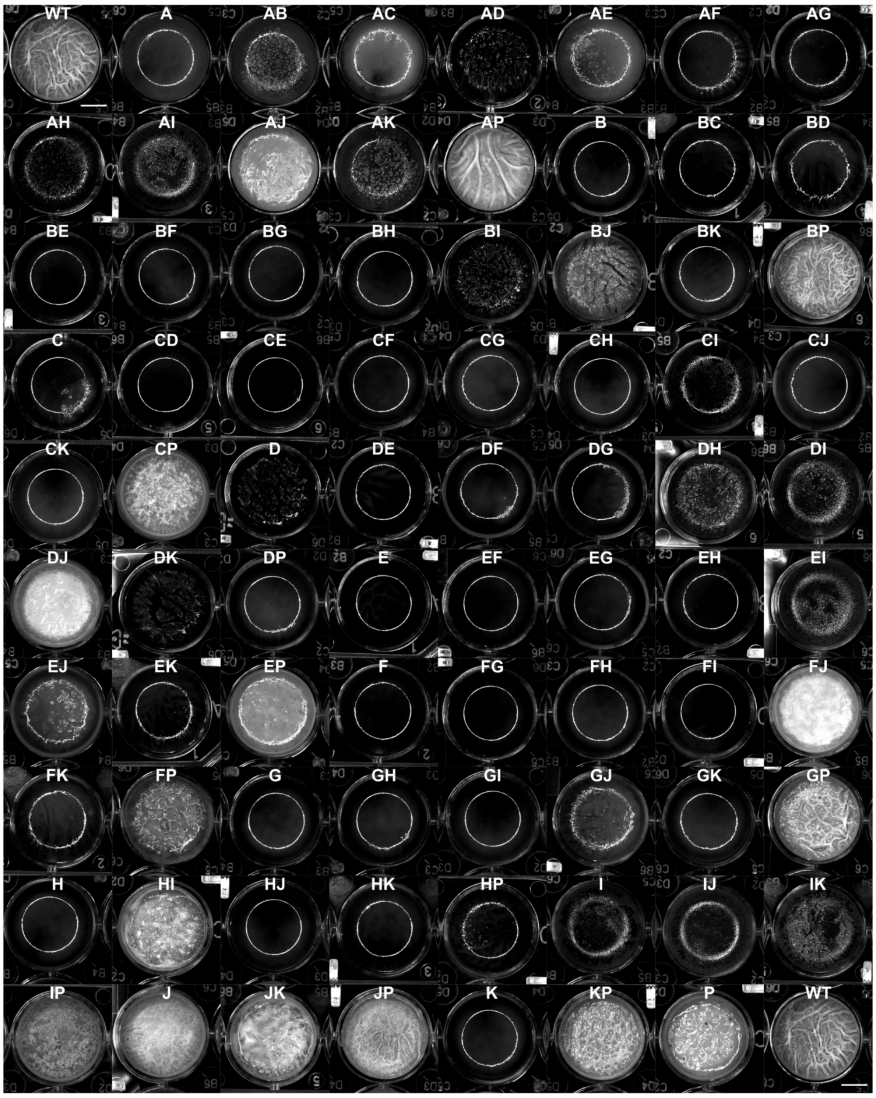


**Supplementary Fig. 4. 5 days pellicles of *B. subtilis* DK1042 and *rap-phr* mutants.** Bright-field images are shown after 5 days of incubation on MSgg medium at 30°C. WT indicates *B. subtilis* DK1042, A indicates a Δ*rapA* mutant, AB indicates a Δ*rapA*Δ*rapB* mutant, and so on. The images shown here have an adjusted contrast, so that the pellicles can be easily appreciated. The same image of *B. subtilis* DK1042 is presented twice (top-left and bottom-right) to facilitate pellicle comparison. The scale bars represent 5 mm.

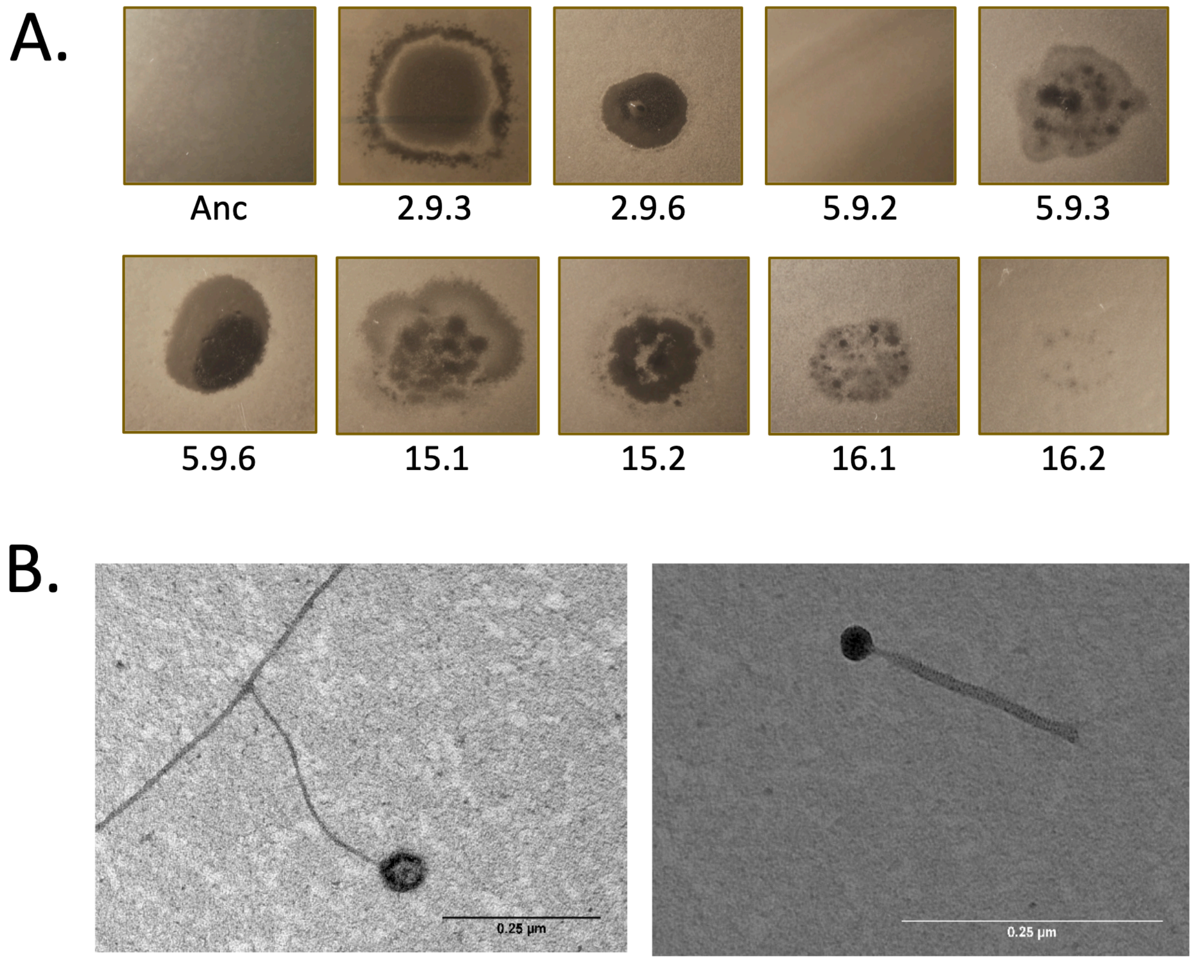


**Supplementary Fig. 5. Presence of infective phage particles in the isolated evolved linages.** (A) Supernatants, obtained from overnight cultures of the evolved strains, were spotted on soft-agar (0.3%) lawn of the ancestor strain with knockout of biofilm genes (NCBI 3610 Δ*eps*Δ*tasA*). (B) Electron microscopy images of isolated phage particles.

**Supplementary Table 1. Strains and plasmids used in this study.**

| **Name** | **Characteristics** | **Reference** |
|---|---|---|
| *B. subtilis* | | |
| NCIB 3610 | Prototroph, wild-type | BGSC |
| DK1042 | NCIB 3610 *comI*$^{Q12I}$ | [1] |
| TB499 | DK1042 *rapA::km*$^{r}$ | This study |
| TB396 | DK1042 *rapC::km*$^{r}$ | This study |
| TB315 | DK1042 *rapD::km*$^{r}$ | This study |
| TB339 | DK1042 *rapE::spec*$^{r}$ | This study |
| TB341 | DK1042 *rapF::spec*$^{r}$ | This study |
| TB404 | DK1042 *rapG::spec*$^{r}$ | This study |
| TB405 | DK1042 *rapH::spec*$^{r}$ | This study |
| TB272 | DK1042 *rapI::km*$^{r}$ | This study |
| TB274 | DK1042 *rapJ::km*$^{r}$ | This study |
| TB557 | DK1042 *rapK::km*$^{r}$ | This study |
| TB435 | DK1042 *rapP::mls*$^{r}$ | This study |
| TB588 | DK1042 Δ*rapA* | This study |
| TB575 | DK1042 Δ*rapB* | This study |
| TB410.1 | DK1042 Δ*rapC* | This study |
| TB513 | DK1042 Δ*rapD* | This study |
| TB407 | DK1042 Δ*rapE* | This study |
| TB408.2 | DK1042 Δ*rapF* | This study |
| TB412 | DK1042 Δ*rapG* | This study |
| TB409.1 | DK1042 Δ*rapH* | This study |
| TB444 | DK1042 Δ*rapI* | This study |
| TB411.2 | DK1042 Δ*rapJ* | This study |
| TB587 | DK1042 Δ*rapK* | This study |
| TB445 | DK1042 Δ*rapP* | This study |
| TB577 | DK1042 Δ*rapB*, *rapA::km*$^{r}$ | This study |
| TB518 | DK1042 Δ*rapC, rapA::km*$^{r}$ | This study |
| TB555 | DK1042 Δ*rapD, rapA::km*$^{r}$ | This study |
| TB517 | DK1042 Δ*rapE, rapA::km*$^{r}$ | This study |
| TB523 | DK1042 Δ*rapF, rapA::km*$^{r}$ | This study |
| TB544 | DK1042 Δ*rapG, rapA::km*$^{r}$ | This study |
| TB522 | DK1042 Δ*rapH, rapA::km*$^{r}$ | This study |
| TB545 | DK1042 Δ*rapI, rapA::km*$^{r}$ | This study |
| TB516 | DK1042 Δ*rapJ, rapA::km*$^{r}$ | This study |
| TB647 | DK1042 Δ*rapK, rapA::km*$^{r}$ | This study |
| TB519 | DK1042 Δ*rapP, rapA::km*$^{r}$ | This study |
| TB582 | DK1042 Δ*rapB, rapC::km*$^{r}$ | This study |
| TB578 | DK1042 Δ*rapB, rapD::km*$^{r}$ | This study |
| TB271 | DK1042 Δ*rapB, rapE::spec*$^{r}$ | This study |
| TB275 | DK1042 Δ*rapB, rapF::spec*$^{r}$ | This study |
| TB583 | DK1042 Δ*rapB, rapG::spec*$^{r}$ | This study |
| TB276 | DK1042 Δ*rapB, rapH::spec*$^{r}$ | This study |
| TB727 | DK1042 Δ*rapB, rapI::km*$^{r}$ | This study |
| TB586 | DK1042 Δ*rapB, rapJ::km*$^{r}$ | This study |
| TB579 | DK1042 Δ*rapB, rapK::km*$^{r}$ | This study |

| | | |
|---|---|---|
| TB584 | DK1042 Δ*rapB, rapP::mls*$^r$ | This study |
| TB521 | DK1042 Δ*rapC, rapD::km*$^r$ | This study |
| TB542 | DK1042 Δ*rapC, rapE::spec*$^r$ | This study |
| TB454 | DK1042 Δ*rapC, rapF::spec*$^r$ | This study |
| TB436 | DK1042 Δ*rapG, rapC::km*$^r$ | This study |
| TB453 | DK1042 Δ*rapC, rapH::spec*$^r$ | This study |
| TB548 | DK1042 Δ*rapI, rapC::km*$^r$ | This study |
| TB455 | DK1042 Δ*rapC, rapJ::km*$^r$ | This study |
| TB561 | DK1042 Δ*rapC, rapK::km*$^r$ | This study |
| TB547 | DK1042 Δ*rapP, rapC::km*$^r$ | This study |
| TB503 | DK1042 Δ*rapE, rapD::km*$^r$ | This study |
| TB504 | DK1042 Δ*rapF, rapD::km*$^r$ | This study |
| TB546 | DK1042 Δ*rapG, rapD::km*$^r$ | This study |
| TB520 | DK1042 Δ*rapH, rapD::km*$^r$ | This study |
| TB550 | DK1042 Δ*rapI, rapD::km*$^r$ | This study |
| TB502 | DK1042 Δ*rapJ, rapD::km*$^r$ | This study |
| TB564 | DK1042 Δ*rapD, rapK::km*$^r$ | This study |
| TB549 | DK1042 Δ*rapP, rapD::km*$^r$ | This study |
| TB456 | DK1042 Δ*rapE, rapF::spec*$^r$ | This study |
| TB451 | DK1042 Δ*rapG, rapE::spec*$^r$ | This study |
| TB457 | DK1042 Δ*rapE, rapH::spec*$^r$ | This study |
| TB285 | DK1042 Δ*rapI, rapE::spec*$^r$ | This study |
| TB458 | DK1042 Δ*rapE, rapJ::km*$^r$ | This study |
| TB558 | DK1042 Δ*rapE, rapK::km*$^r$ | This study |
| TB283 | DK1042 Δ*rapP, rapE::spec*$^r$ | This study |
| TB452 | DK1042 Δ*rapG, rapF::spec*$^r$ | This study |
| TB543 | DK1042 Δ*rapH, rapF::spec*$^r$ | This study |
| TB472 | DK1042 Δ*rapI, rapF::spec*$^r$ | This study |
| TB495 | DK1042 Δ*rapJ, rapF::spec*$^r$ | This study |
| TB559 | DK1042 Δ*rapF, rapK::km*$^r$ | This study |
| TB293 | DK1042 Δ*rapP, rapF::spec*$^r$ | This study |
| TB433 | DK1042 Δ*rapG, rapH::spec*$^r$ | This study |
| TB474 | DK1042 Δ*rapI, rapG::spec*$^r$ | This study |
| TB434 | DK1042 Δ*rapG, rapJ::km*$^r$ | This study |
| TB566 | DK1042 Δ*rapG, rapK::km*$^r$ | This study |
| TB443 | DK1042 Δ*rapG, rapP::mls*$^r$ | This study |
| TB473 | DK1042 Δ*rapI, rapH::spec*$^r$ | This study |
| TB496 | DK1042 Δ*rapJ, rapH::spec*$^r$ | This study |
| TB560 | DK1042 Δ*rapH, rapK::km*$^r$ | This study |
| TB284 | DK1042 Δ*rapP, rapH::spec*$^r$ | This study |
| TB551 | DK1042 Δ*rapI, rapJ::km*$^r$ | This study |
| TB562 | DK1042 Δ*rapI, rapK::km*$^r$ | This study |
| TB552 | DK1042 Δ*rapI, rapP::mls*$^r$ | This study |
| TB565 | DK1042 Δ*rapJ, rapK::km*$^r$ | This study |
| TB292 | DK1042 Δ*rapJ, rapP::mls*$^r$ | This study |
| TB563 | DK1042 Δ*rapP, rapK::km*$^r$ | This study |
| TB614.BC | DK1042 *amyE::*barcode (*cat*$^r$) | This study |
| *E. coli* | | |

| | | |
|---|---|---|
| MC1061 | Cloning host; K-12 $F^{-}$ $\lambda^{-}$ $\Delta$(*ara-leu*)7697 [*araD139*]B/r $\Delta$(*codB-lacI*)3 *galK16 galE15 e14*$^{-}$ *mcrA0 relA1 rpsL150*(*Str*$^{r}$) *spoT1 mcrB1 hsdR2*($r^{-}m^{+}$) | [2] |
| **Plasmid** | **Characteristics** | **Reference** |
| pMAD_rapB | pMAD thermosensitive plasmid, *rapB*-5´, *rapB*-3´ | provided by Stephanie Trauth (Ilka Bischofs´ Laboratory) |
| pBluescript Sk(+) | Cloning vector. *Amp*$^{r}$, *lacZ* | Stratagene |
| pTB120 | pBluescript Sk(+) *lacZ::lox66-neo*$^{r}$*-lox71* | [3] |
| pTB233 | pBluescript Sk(+) *lacZ::lox66-spec*$^{r}$*-lox71* | This study |
| pTB234 | pBluescript Sk(+) *lacZ::lox66-mls*$^{r}$*-lox71* | This study |
| pTB250 | pTB120 *rapI*-5´*-lox66-neo*$^{r}$*-lox71-phrI*-3´ | This study |
| pTB251 | pTB120 *rapJ*-5´*-lox66-neo*$^{r}$*-lox71-rapJ*-3´ | This study |
| pTB252 | pTB120 *rapK*-5´*-lox66-neo*$^{r}$*-lox71-phrK*-3´ | This study |
| pTB295 | pTB120 *rapD*-5´*-lox66-neo*$^{r}$*-lox71-rapD*-3´ | This study |
| pTB310 | pTB233 *rapE*-5´*-lox66-spec*$^{r}$*-lox71-phrE*-3´ | This study |
| pTB311 | pTB233 *rapF*-5´*-lox66-spec*$^{r}$*-lox71-phrF*-3´ | This study |
| pTB349 | pTB233 *rapH*-5´*-lox66-spec*$^{r}$*-lox71-phrH*-3´ | This study |
| pTB380 | pTB120 *rapA*-5´*-lox66-neo*$^{r}$*-lox71-phrA*-3´ | This study |
| pTB382 | pTB120 *rapC*-5´*-lox66-neo*$^{r}$*-lox71-phrC*-3´ | This study |
| pTB383 | pTB233 *rapG*-5´*-lox66-spec*$^{r}$*-lox71-phrG*-3´ | This study |
| pTB414 | pTB234 *rapP*-5´*-lox66-mls*$^{r}$*-lox71-phrP*-3´ | This study |
| pMH66 | pNZ124-based Cre-encoding plasmid, *Tet*$^{r}$ *Ts* | [4] |
| pTB16 | pDG782 derivate plasmid. *amyE* integration vector for *B. subtilis. km*$^{r}$ | [5] |
| pNW33n | *cat*$^{r}$, *Geobacillus-E. coli* shuttle vector | BGSC |
| pTB666.1 to pTB666.80 | pTB16 derivate plasmids. *amyE*-5´*-cat*$^{r}$ (barcode)-*amyE*-3´. The barcodes are 12-bp random nucleotide sequences | This study |

Note: All *B. subtilis* strains with at least one deletion of a *rap-phr* pair were tagged with a barcode using a pTB666 plasmid (see TB614.BC) and listed in S4 Table. The resulting barcoded strains have no additional modifications compared to their parental strain.

**Supplementary Table 2. Primers used in this study**

| Primer | Target locus | Sequence (5´→3´) |
|---|---|---|
| oRGM2 | *neo-lox66* | TACCGTTCGTATAATGTATGCTATACGAAGTTATAGATCAATTTGATA ATTACTAATAC |
| oRGM7 | *neo-lox71* | TACCGTTCGTATAGCATACATTATACGAAGTTATTAGAGCTTGGGTT ACAGGCATGG |
| oRGM14 | *mls-lox71* | TACCGTTCGTATAGCATACATTATACGAAGTTATAGAAACGCAAAAA GGCCATCCGTCAG |
| oRGM15 | *mls-lox66* | TACCGTTCGTATAATGTATGCTATACGAAGTTATCCTACCGCGGGC GGCCGCACTCTTCC |
| oRGM16 | *spe-lox71* | TACCGTTCGTATAGCATACATTATACGAAGTTATCTCGAGATCCCCC TATGCAAGGG |
| oRGM17 | *spe-lox66* | TACCGTTCGTATAATGTATGCTATACGAAGTTATTAAAAAATTTAGAA GCCAATGAAATC |
| oRGM20 | *rapI* 5´ | ATCCTCGAGTGGTTCCTCCAAGGAGAATG |
| oRGM21 | *rapI* 5´ | ACGCTGCAGGTGACTAAGTCGTACGG |
| oRGM22 | *rapI* 3´ | ATCGGATCCAGTTGCTGCAGATCGGGTAG |
| oRGM23 | *rapI* 3´ | ACGGAGCTCACCATTGTTTGGTCGTTCTG |
| oRGM24 | *rapI* | TTGGTGCTACTAGCAGTG |
| oRGM25 | *rapI* | GGGCAGCAAACTCATAGTTC |
| oRGM26 | *rapJ* 5´ | ATCGGTACCTATGCCCTCTATCCGAGAGC |
| oRGM27 | *rapJ* 5´ | ACGGAATTCTGCGCGAATGAGCTTGTACC |
| oRGM28 | *rapJ* 3´ | ATCGGATCCAAAGAAGCTTGCCGAGCAG |
| oRGM29 | *rapJ* 3´ | ACGGAGCTCGTCAAGACGGGAAATAATC |
| oRGM30 | *rapJ* | CCTCCAATGCTCCACGGAAG |
| oRGM31 | *rapJ* | GGATAGATCGGGCAAATCC |
| oRGM32 | *rapK* 5´ | ATCGGTACCTCTTCTGTTACCGCTGAGTC |
| oRGM33 | *rapK* 5´ | ACGGAATTCAACTTCAGAAGCGATCTTAC |
| oRGM34 | *rapK* 3´ | ATCGGATCCACATCCAGGTAGCTGAAAGG |
| oRGM35 | *rapK* 3´ | ATGCGGCCGCAAACAGGATCGAGACTATTTG |
| oRGM36 | *rapK* | GCGGTCTTTTATGTATGAAATC |
| oRGM37 | *rapK* | GGATAGACAGGGAAGTGTAG |
| oRGM44 | *rapD* 3´ | GGATCCAAAAGCCGCTTTTTTTATCATG |
| oRGM45 | *rapD* 3´ | ACGGAGCTCTGACTGAAGCGTACAGATCG |
| oRGM46 | *rapD* | TTGCTGCTTCAGCAGGTCTC |
| oRGM47 | *rapD* | GCGTCTCAGAGCTTTCAAAC |
| oRGM48 | *rapE* 5´ | ATGGGCCCGGCCAATCAGCTGGATCTTC |
| oRGM49 | *rapE* 5´ | AGCTGCAGATCTTCATTCCCACTTCAG |
| oRGM50 | *rapE* 3´ | ATGGATCCTGTAACTCTCGCACCTACTC |
| oRGM51 | *rapE* 3´ | ACGAGCTCATGTTATTAGCGCCTTTGCC |
| oRGM52 | *rapE* | TTTGCTGTGAGCCGGTGTAG |
| oRGM53 | *rapE* | GCAATGCCAGCTTGATCTTC |
| oRGM54 | *rapF* 5´ | ATGGGCCCGATTCGCTGTAAACGCGTAG |
| oRGM55 | *rapF* 5´ | CTGAATTCGTATGCTGAATCGGCGTATG |
| oRGM56 | *rapF* 3´ | ATGGATCCGAAGTTGCACAACGAGGAATG |
| oRGM57 | *rapF* 3´ | ACGAGCTCCGGCGGCATCACGTCTAAAG |
| oRGM58 | *rapF* | ACGGAAGAGCAATCGTTGTC |
| oRGM59 | *rapF* | GGCCGTCCGGTTTATGTCAC |

| | | |
|---|---|---|
| oRGM62 | *rapG* 3´ | ATGGATCCCGGACCATCAAACCCACTCAC |
| oRGM63 | *rapG* 3´ | ATGCGGCCGCACGGCGATTTGAATACACTTG |
| oRGM64 | *rapG* | TGCAGTGCGGCGATTTCTTC |
| oRGM65 | *rapG* | TATTGCGATCGGCACGCTTG |
| oRGM66 | *rapH* 5´ | ATGGGCCCTTGATACGACGGGAAATGAG |
| oRGM67 | *rapH* 5´ | CTGCAGCGCGAAGACGGTATGGCTTGAC |
| oRGM68 | *rapH* 3´ | GGATCCATTCCCCTTACAAACTTAGTG |
| oRGM69 | *rapH* 3´ | GCTCTAGAATCCGGAAGCGTTACTTCAC |
| oRGM70 | *rapH* | CCGCTGTCAGATCCATTTGC |
| oRGM71 | *rapH* | CCTGCTCACTCCTTACTCAC |
| oRGM72 | *rapA* 5´ | ATGGTACCCAGTATCGATGCACCTGTTG |
| oRGM73 | *rapA* 5´ | ATGAATTCCGGCTTCAGCGACGTGGAAC |
| oRGM74 | *rapA* 3´ | GTTCTAGATGCGGCACGCAATCAAAC |
| oRGM75 | *rapA* 3´ | CTGAGCTCAGGCTTCAGCTGCCTCATAC |
| oRGM76 | *rapA* | CGCGGCATTCTGTTATATGG |
| oRGM77 | *rapA* | TCCAGTCCTGATGCTTTCTC |
| oRGM84 | *rapC* 5´ | ATGGTACCGACGACGATCAACGGTTTGG |
| oRGM85 | *rapC* 5´ | ATCTGCAGTTGACCGACCGCTGAAGAAG |
| oRGM86 | *rapC* 3´ | ATGGATCCCTAATGCGGAAGCACTCGAC |
| oRGM87 | *rapC* 3´ | ATGAGCTCGGATTTGCATGCCGATGAAG |
| oRGM88 | *rapC* | AATCGAGCGCCTTGAGAAGC |
| oRGM89 | *rapC* | TCGGGAATCGATGACATGAC |
| oRGM90 | *rapD* 5´ | ATGGTACCTTCCGAAAGCGCCGCCTATC |
| oRGM91 | *rapD* 5´ | ATCTGCAGCGGAATACCACTCGTCTAAC |
| oRGM92 | *rapG* 5´ | ATGTCGACCGCACATTGTGAGCGCTACC |
| oRGM93 | *rapG* 5´ | ATCTGCAGTGATGGCAAGGTACCAATCG |
| oRGM94 | *rapP* 5´ | ATGGGCCCTCCCAATCGTTTGGAGAAAG |
| oRGM95 | *rapP* 5´ | ATGAATTCTGGGATTAAATCCGAAAC |
| oRGM96 | *rapP* 3´ | ATGGATCCACTTATAAGGTCGCAGATAG |
| oRGM97 | *rapP* 3´ | ATGAGCTCGGGCTGCATATAAATAATAAG |
| oRGM98 | *rapP* | TCCAACGTGCAGTGGAAAGG |
| oRGM99 | *rapP* | CTTCACTCAAGAAGAACAAG |
| oTB118 | *Cm*$^{R}$ | GATCAGATCTCCGGCGTAGAGGATCTGG |
| oTB119 | *Cm*$^{R}$ (barcode) | CACGAAGCTTGCNNNNNNNNNNNNTATCATCGGCAATAGTTACCC |
| oTB120 | *amyE* | GAGGAAGCGGAAGAATGAAG |
| oTB121 | *amyE* | TTCGGTAAGTCCCGTCTAGC |
| oBC_fw | barcode locus | TGCGGTGATTGTTAGGTTGAGGCCGTTGAG |
| oBC_rev | barcode locus | AGTCAGTCAGCCAGGAGGCTTACTTGTCTG |
| oBC1 | barcode locus (fw) | AATGATACGGCGACCACCGAGATCTACACATCGTACGTGCGGTGAT TGTTAGGTTGAGGCCGTTGAG |
| oBC2 | barcode locus (fw) | AATGATACGGCGACCACCGAGATCTACACACTATCTGTGCGGTGAT TGTTAGGTTGAGGCCGTTGAG |
| oBC3 | barcode locus (fw) | AATGATACGGCGACCACCGAGATCTACACTAGCGAGTTGCGGTGAT TGTTAGGTTGAGGCCGTTGAG |
| oBC4 | barcode locus (fw) | AATGATACGGCGACCACCGAGATCTACACCTGCGTGTTGCGGTGAT TGTTAGGTTGAGGCCGTTGAG |
| oBC5 | barcode locus (rv) | CAAGCAGAAGACGGCATACGAGATAACTCTCGAGTCAGTCAGCCA GGAGGCTTACTTGTCTG |
| oBC6 | barcode locus (rv) | CAAGCAGAAGACGGCATACGAGATACTATGTCAGTCAGTCAGCCAG |

| | | |
|---|---|---|
| | | GAGGCTTACTTGTCTG |
| oBC7 | barcode locus (rv) | CAAGCAGAAGACGGCATACGAGATAGTAGCGTAGTCAGTCAGCCAGGAGGCTTACTTGTCTG |
| oBC8 | barcode locus (rv) | CAAGCAGAAGACGGCATACGAGATCAGTGAGTAGTCAGTCAGCCAGGAGGCTTACTTGTCTG |
| oBC9 | barcode locus (rv) | CAAGCAGAAGACGGCATACGAGATCGTACTCAAGTCAGTCAGCCAGGAGGCTTACTTGTCTG |
| oBC10 | barcode locus (rv) | CAAGCAGAAGACGGCATACGAGATCTACGCAGAGTCAGTCAGCCAGGAGGCTTACTTGTCTG |
| oBC11 | barcode locus (rv) | CAAGCAGAAGACGGCATACGAGATGGAGACTAAGTCAGTCAGCCAGGAGGCTTACTTGTCTG |
| oBC12 | barcode locus (rv) | CAAGCAGAAGACGGCATACGAGATGTCGCTCGAGTCAGTCAGCCAGGAGGCTTACTTGTCTG |
| oBC13 | barcode locus (rv) | CAAGCAGAAGACGGCATACGAGATGTCGTAGTAGTCAGTCAGCCAGGAGGCTTACTTGTCTG |
| oBC14 | barcode locus (rv) | CAAGCAGAAGACGGCATACGAGATTAGCAGACAGTCAGTCAGCCAGGAGGCTTACTTGTCTG |
| oBC15 | barcode locus (rv) | CAAGCAGAAGACGGCATACGAGATTCATAGACAGTCAGTCAGCCAGGAGGCTTACTTGTCTG |
| oBC16 | barcode locus (rv) | CAAGCAGAAGACGGCATACGAGATTCGCTATAAGTCAGTCAGCCAGGAGGCTTACTTGTCTG |

**Supplementary Table 3. DNA barcoded strains used in this study.**

| Name | Characteristics | DNA barcode |
|---|---|---|
| TB614.BC | DK1042 *amyE::*barcode (*cat*$^{r}$) | CCCTAATGAGAA |
| TB588.BC | DK1042 Δ*rapA amyE::*barcode (*cat*$^{r}$) | TTGGCCATTGTG |
| TB575.BC | DK1042 Δ*rapB amyE::*barcode (*cat*$^{r}$) | TCTTCTGGAGCC |
| TB410.1.BC | DK1042 Δ*rapC amyE::*barcode (*cat*$^{r}$) | AATTTCGAGTCG |
| TB513.BC | DK1042 Δ*rapD amyE::*barcode (*cat*$^{r}$) | ATTGCTTTTTTT |
| TB407.BC | DK1042 Δ*rapE amyE::*barcode (*cat*$^{r}$) | GGTAGGGCATTG |
| TB408.2.BC | DK1042 Δ*rapF amyE::*barcode (*cat*$^{r}$) | CAGGGGTTGCAC |
| TB412.BC | DK1042 Δ*rapG amyE::*barcode (*ca*$^{r}$) | GCATGCGAGTCG |
| TB409.1.BC | DK1042 Δ*rapH amyE::*barcode (*cat*$^{r}$) | TCGGGTGATAGT |
| TB444.BC | DK1042 Δ*rapI amyE::*barcode (*cat*$^{r}$) | TTGTTGAACACC |
| TB411.2.BC | DK1042 Δ*rapJ amyE::*barcode (*cat*$^{r}$) | GGAAGGATTATG |
| TB587.BC | DK1042 Δ*rapK amyE::*barcode (*cat*$^{r}$) | GGGTTACATATT |
| TB445.BC | DK1042 Δ*rapP amyE::*barcode (*cat*$^{r}$) | TATTTCATGGAT |
| TB577.BC | DK1042 Δ*rapB*, *rapA::km*$^{R}$ *amyE::*barcode (*cat*$^{r}$) | GGAACGGGTCGT |
| TB518.BC | DK1042 Δ*rapC, rapA::km*$^{R}$ *amyE::*barcode (*cat*$^{r}$) | GGTGGGTGTGAG |
| TB555.BC | DK1042 Δ*rapD, rapA::km*$^{R}$ *amyE::*barcode (*cat*$^{r}$) | GGTAGGGGCCAG |
| TB517.BC | DK1042 Δ*rapE, rapA::km*$^{R}$ *amyE::*barcode (*cat*$^{r}$) | GATTGAGCCAGC |
| TB523.BC | DK1042 Δ*rapF, rapA::km*$^{R}$ *amyE::*barcode (*cat*$^{r}$) | CTGATACCGTTT |
| TB544.BC | DK1042 Δ*rapG, rapA::km*$^{R}$ *amyE::*barcode (*cat*$^{r}$) | GGCTCCGTTTAG |
| TB522.BC | DK1042 Δ*rapH, rapA::km*$^{R}$ *amyE::*barcode (*cat*$^{r}$) | TCATCTTCTGGT |
| TB545.BC | DK1042 Δ*rapI, rapA::km*$^{R}$ *amyE::*barcode (*cat*$^{r}$) | GACGGTCGGTGT |
| TB516.BC | DK1042 Δ*rapJ, rapA::km*$^{R}$ *amyE::*barcode (*cat*$^{r}$) | TTTAGTTTGGAC |
| TB647.BC | DK1042 Δ*rapK, rapA::km*$^{R}$ *amyE::*barcode (*cat*$^{r}$) | AGTCGTTGTACG |
| TB519.BC | DK1042 Δ*rapP, rapA::km*$^{R}$ *amyE::*barcode (*cat*$^{r}$) | ATTAGTTGTTAC |
| TB582.BC | DK1042 Δ*rapB, rapC::km*$^{R}$ *amyE::*barcode (*cat*$^{r}$) | GTCTTGGGGAGG |
| TB578.BC | DK1042 Δ*rapB, rapD::km*$^{R}$ *amyE::*barcode (*cat*$^{r}$) | TTTGGGGCCCGG |
| TB271.BC | DK1042 Δ*rapB, rapE::spec*$^{R}$ *amyE::*barcode (*cat*$^{r}$) | TCCGGGAATGAA |
| TB275.BC | DK1042 Δ*rapB, rapF::spec*$^{R}$ *amyE::*barcode (*cat*$^{r}$) | GGTTGTTCCTCT |
| TB583.BC | DK1042 Δ*rapB, rapG::spec*$^{R}$ *amyE::*barcode (*cat*$^{r}$) | GGGGGGGTGTTT |
| TB276.BC | DK1042 Δ*rapB, rapH::spec*$^{R}$ *amyE::*barcode (*cat*$^{r}$) | TCACAGACATTG |
| TB727.BC | DK1042 Δ*rapB, rapI::km*$^{R}$ *amyE::*barcode (*cat*$^{r}$) | CACATGACCAGA |
| TB586.BC | DK1042 Δ*rapB, rapJ::km*$^{R}$ *amyE::*barcode (*cat*$^{r}$) | GGTAGCTGGTCC |
| TB579.BC | DK1042 Δ*rapB, rapK::km*$^{R}$ *amyE::*barcode (*cat*$^{r}$) | TATAGGCCTTCG |
| TB584.BC | DK1042 Δ*rapB, rapP::mls*$^{R}$ *amyE::*barcode (*cat*$^{r}$) | AACACAAAGTAC |
| TB521.BC | DK1042 Δ*rapC, rapD::km*$^{R}$ *amyE::*barcode (*cat*$^{r}$) | GGGTTGTAGTGC |
| TB542.BC | DK1042 Δ*rapC, rapE::spec*$^{R}$ *amyE::*barcode (*cat*$^{r}$) | GCGCTAGTCCTA |
| TB454.BC | DK1042 Δ*rapC, rapF::spec*$^{R}$ *amyE::*barcode (*cat*$^{r}$) | GGTAGAGCTGTC |
| TB436.BC | DK1042 Δ*rapG, rapC::km*$^{R}$ *amyE::*barcode (*cat*$^{r}$) | GCGTAAGGGTAG |
| TB453.BC | DK1042 Δ*rapC, rapH::spec*$^{R}$ *amyE::*barcode (*cat*$^{r}$) | CTCTGCAACAAT |
| TB548.BC | DK1042 Δ*rapI, rapC::km*$^{R}$ *amyE::*barcode (*cat*$^{r}$) | GACACCCCCATC |
| TB455.BC | DK1042 Δ*rapC, rapJ::km*$^{R}$ *amyE::*barcode (*cat*$^{r}$) | TCAGTGAGGATG |
| TB561.BC | DK1042 Δ*rapC, rapK::km*$^{R}$ *amyE::*barcode (*cat*$^{r}$) | ATCGGAGTGGAG |
| TB547.BC | DK1042 Δ*rapP, rapC::km*$^{R}$ *amyE::*barcode (*cat*$^{r}$) | TATGGGCTGACG |
| TB503.BC | DK1042 Δ*rapE, rapD::km*$^{R}$ *amyE::*barcode (*cat*$^{r}$) | CGGCGTCTCGGG |
| TB504.BC | DK1042 Δ*rapF, rapD::km*$^{R}$ *amyE::*barcode (*cat*$^{r}$) | TTGAGCGCGGTG |
| TB546.BC | DK1042 Δ*rapG, rapD::km*$^{R}$ *amyE::*barcode (*cat*$^{r}$) | GGGAGAGGCAGG |

| | | |
|---|---|---|
| TB520.BC | DK1042 Δ*rapH, rapD::km*$^{R}$ *amyE::*barcode (*cat*$^{r}$) | GGCAGGCTTGTA |
| TB550.BC | DK1042 Δ*rapI, rapD::km*$^{R}$ *amyE::*barcode (*cat*$^{r}$) | GGTATCGAAGGC |
| TB502.BC | DK1042 Δ*rapJ, rapD::km*$^{R}$ *amyE::*barcode (*cat*$^{r}$) | TAAGTCCGTTAG |
| TB564.BC | DK1042 Δ*rapD, rapK::km*$^{R}$ *amyE::*barcode (*cat*$^{r}$) | GCGTGATCCGGT |
| TB549.BC | DK1042 Δ*rapP, rapD::km*$^{R}$ *amyE::*barcode (*cat*$^{r}$) | GTTTTCCAATGC |
| TB456.BC | DK1042 Δ*rapE, rapF::spec*$^{R}$ *amyE::*barcode (*cat*$^{r}$) | GCTGACGGGGAA |
| TB451.BC | DK1042 Δ*rapG, rapE::spec*$^{R}$ *amyE::*barcode (*cat*$^{r}$) | TGTAGCGCTGGT |
| TB457.BC | DK1042 Δ*rapE, rapH::spec*$^{R}$ *amyE::*barcode (*cat*$^{r}$) | GACTTTAAGATG |
| TB285.BC | DK1042 Δ*rapI, rapE::spec*$^{R}$ *amyE::*barcode (*cat*$^{r}$) | TCTTTAGTGAAA |
| TB458.BC | DK1042 Δ*rapE, rapJ::km*$^{R}$ *amyE::*barcode (*cat*$^{r}$) | GGATTCCAAACG |
| TB558.BC | DK1042 Δ*rapE, rapK::km*$^{R}$ *amyE::*barcode (*cat*$^{r}$) | GCGGACGTCATG |
| TB283.BC | DK1042 Δ*rapP, rapE::spec*$^{R}$ *amyE::*barcode (*cat*$^{r}$) | AATGGTCAACTG |
| TB452.BC | DK1042 Δ*rapG, rapF::spec*$^{R}$ *amyE::*barcode (*cat*$^{r}$) | CCGGTTATGGCG |
| TB543.BC | DK1042 Δ*rapH, rapF::spec*$^{R}$ *amyE::*barcode (*cat*$^{r}$) | TGGGTGTTTGAT |
| TB472.BC | DK1042 Δ*rapI, rapF::spec*$^{R}$ *amyE::*barcode (*cat*$^{r}$) | TGTTTGTGCTTT |
| TB495.BC | DK1042 Δ*rapJ, rapF::spec*$^{R}$ *amyE::*barcode (*cat*$^{r}$) | GGTTAGGGTCCC |
| TB559.BC | DK1042 Δ*rapF, rapK::km*$^{R}$ *amyE::*barcode (*cat*$^{r}$) | CAATAATCGCTT |
| TB293.BC | DK1042 Δ*rapP, rapF::spec*$^{R}$ *amyE::*barcode (*cat*$^{r}$) | GTCACGCACGTT |
| TB433.BC | DK1042 Δ*rapG, rapH::spec*$^{R}$ *amyE::*barcode (*cat*$^{r}$) | GGGACGGTTTGT |
| TB474.BC | DK1042 Δ*rapI, rapG::spec*$^{R}$ *amyE::*barcode (*cat*$^{r}$) | GGGCAGCTGAGA |
| TB434.BC | DK1042 Δ*rapG, rapJ::km*$^{R}$ *amyE::*barcode (*cat*$^{r}$) | GTTACTTTAGCG |
| TB566.BC | DK1042 Δ*rapG, rapK::km*$^{R}$ *amyE::*barcode (*cat*$^{r}$) | TTGAGTCTTCGC |
| TB443.BC | DK1042 Δ*rapG, rapP::mls*$^{R}$ *amyE::*barcode (*cat*$^{r}$) | GTGTTATAGTAT |
| TB473.BC | DK1042 Δ*rapI, rapH::spec*$^{R}$ *amyE::*barcode (*cat*$^{r}$) | TCGCTAATTACA |
| TB496.BC | DK1042 Δ*rapJ, rapH::spec*$^{R}$ *amyE::*barcode (*cat*$^{r}$) | GTCAACAGATAT |
| TB560.BC | DK1042 Δ*rapH, rapK::km*$^{R}$ *amyE::*barcode (*cat*$^{r}$) | TTTAGTCCGGAA |
| TB284.BC | DK1042 Δ*rapP, rapH::spec*$^{R}$ *amyE::*barcode (*cat*$^{r}$) | GGTTGGGCGGGT |
| TB551.BC | DK1042 Δ*rapI, rapJ::km*$^{R}$ *amyE::*barcode (*cat*$^{r}$) | CCAAGGCCCGTG |
| TB562.BC | DK1042 Δ*rapI, rapK::km*$^{R}$ *amyE::*barcode (*cat*$^{r}$) | GCTCGAATCCCG |
| TB552.BC | DK1042 Δ*rapI, rapP::mls*$^{R}$ *amyE::*barcode (*cat*$^{r}$) | TGCCACGGAAAG |
| TB565.BC | DK1042 Δ*rapJ, rapK::km*$^{R}$ *amyE::*barcode (*cat*$^{r}$) | CTTGTTGAAACA |
| TB292.BC | DK1042 Δ*rapJ, rapP::mls*$^{R}$ *amyE::*barcode (*cat*$^{r}$) | GCATGCACCGTA |
| TB563.BC | DK1042 Δ*rapP, rapK::km*$^{R}$ *amyE::*barcode (*cat*$^{r}$) | TGGGCTGTGGCC |

**Supplementary Dataset 1. List of mutations detected in genomes of sequenced population clones. Mutations detected in each sequenced clone are indicated in separate tabs, while the last two tabs enlist mutations found in multiple and in single evolved isolates, respectively.**
